# Proximity mapping of desmosomes reveals a striking shift in their molecular neighbourhood associated with maturation

**DOI:** 10.1101/2023.03.24.534085

**Authors:** Judith B. Fülle, Rogerio Alves de Almeida, Craig Lawless, Bian Yanes, E. Birgitte Lane, David R. Garrod, Christoph Ballestrem

**Author notes:** These authors contributed equally to this work (Judith B. Fülle, Rogerio Alves de Almeida).

## Abstract

Desmosomes are multiprotein adhesion complexes that link intermediate filaments to the plasma membrane, ensuring the mechanical integrity of cells across tissues, but how they participate in the wider signalling network to exert their full function is unclear. To investigate this we carried out multiplexed protein proximity mapping using biotinylation (BioID). The combined interactomes of the essential desmosomal proteins desmocollin 2a, plakoglobin and plakophilin 2a (Pkp2a) in Madin-Darby canine kidney epithelial cells were mapped and their differences and commonalities characterised as desmosome matured from Ca^2+^-dependence to the mature, Ca^2+^-independent, hyperadhesive state, which predominates in tissues. Results suggest that individual desmosomal proteins have distinct roles in connecting to cellular signalling pathways and that these roles alter substantially when cells change their adhesion state. The data provide further support for a dualistic concept of desmosomes in which the properties of Pkp2a differ from those of the other, more stable proteins. This body of data provides an invaluable resource for analysis of desmosome function.

## Introduction

Desmosomes are complex, multiprotein cell-cell junctions providing strong intercellular adhesion and connecting to cytoplasmic intermediate filament (IF) networks. These functions are especially vital for mechanically resilient tissues such as the epidermis and cardiac muscle. The association of impaired desmosome function with severe diseases has revealed that both mechanical resilience and multiple signalling pathways are disrupted in affected tissues (Green et al., 2019; Müller et al., 2021; Spindler et al., 2018). However, the underlying mechanisms are poorly understood.

The canonical desmosomal complex comprises two kinds of cell adhesion receptors (the desmosomal cadherin proteins desmogleins (Dsg1-4) and desmocollins (Dsc1-3 a and b)), two adaptor proteins (the armadillo proteins plakophilins (Pkp1-3) and plakoglobin (PG, gene name JUP)) and the plakin protein desmoplakin (DP). Desmoplakin connects the desmosome junction to the IF network of the cell, forming the desmosome-IF complex, a mechanically resilient tissue-wide scaffolding (reviewed by Delva et al., 2009; Garrod and Chidgey, 2008; Green et al., 2019).

During and shortly after assembly in tissue culture, desmosomal adhesion is Ca^2+^-dependent and can be disrupted by removal of extracellular calcium. In vivo, this default state is seen in early embryonic tissues and wound epidermis (Berika and Garrod, 2014). Following assembly, Ca^2+^-dependent desmosomes mature, both in culture and in vivo, to adopt a more stable Ca^2+^-independent adhesion state referred to as hyper-adhesion which is the normal condition found in epithelia at homeostasis (Berika and Garrod, 2014; Kimura et al., 2012; Wallis et al., 2000). This maturation process appears characteristic of desmosomes and distinguishes them from adherens junctions (AJ), the other major type of adhesive epithelial junction. How cells regulate desmosomal adhesion and what consequences the change of adhesion state signify is not understood.

Desmosomes are remarkably stable structures. They persist even when cells are undergoing mitosis (Baker and Garrod, 1993), and when epithelial cells separate, desmosomes are torn out of the neighbouring cell to be engulfed as whole entities (Fülle et al., 2021). However, overall stability is only part of the story, because the desmosomal complex exhibits dualistic properties, as we recently showed (Fülle et al., 2021). Thus, while the DCs, PG and DP remain quite stably associated with the junctions, at least one protein, Pkp2a, is remarkably dynamic, shuttling in and out of desmosomes to other cellular compartments within seconds. This behaviour of Pkp2a together with the presence of PG and Pkps in other cellular compartments, including the nucleus, supports a model whereby desmosomes are not simply mediating firm adhesion but also function as signalling platforms (Müller et al., 2021). Very little is so far known about how such signalling is organised and how desmosomes connect to signalling networks. It is also unclear how the adhesion state of desmosomes, which is often neglected despite its importance, is regulated and how it affects signalling capacities and pathways.

Here, we have used BioID-coupled mass spectrometry to study how desmosomal components are embedded in signalling networks (Roux et al., 2018). To reveal changes in relation to desmosome adhesion states we generated proximitomes from Dsc2a, PG and Pkp2a baits with either Ca^2+^-dependent or hyper-adhesive desmosomes. The resulting combined network showed substantial changes in proximal proteins and their possible functions as desmosomes mature, as well as revealing novel partners with potentially novel signalling roles. Moreover, the results reinforce our view of the dualistic nature of desmosomal protein for while Dsc2a and PG share many partners with common functions those of Pkp2a differ considerably. The results expand our view of desmosomes and their roles beyond mechanically resilient cell-cell junctions.

## Results and Discussion

### Proximitome of core desmosome components

To investigate the proximitome of desmosomes, multiplexed BioID was carried out (Fig.1A). We selected Dsc2a, PG and Pkp2a as BioID baits. The proteins were cloned into a custom-made vector containing BirA-myc and stably expressed in the simple epithelial cell line Madin-Darby canine kidney type II (MDCK). Dsc2a was tagged on its cytoplasmic C-terminus (Dsc2a-BirA-myc, Dsc2a*), PG on both the N- (myc-BirA-PG, PG-N*) and C-termini (PG-BirA-myc, PG-C*), and Pkp2a on its N-terminus (myc-BirA-Pkp2a, Pkp2a*). Tagging of the C-terminus of Pkp2a was dismissed as it interfered with localisation to desmosomes (data not shown), confirming findings that the C-terminus of Pkps, in particular of Pkp1, is required for recruitment to the plasma membrane (Sobolik-Delmaire et al., 2006). As controls, we generated MDCK cells stably expressing BirA-myc alone (Roux et al., 2018).

**Fig. 1.**
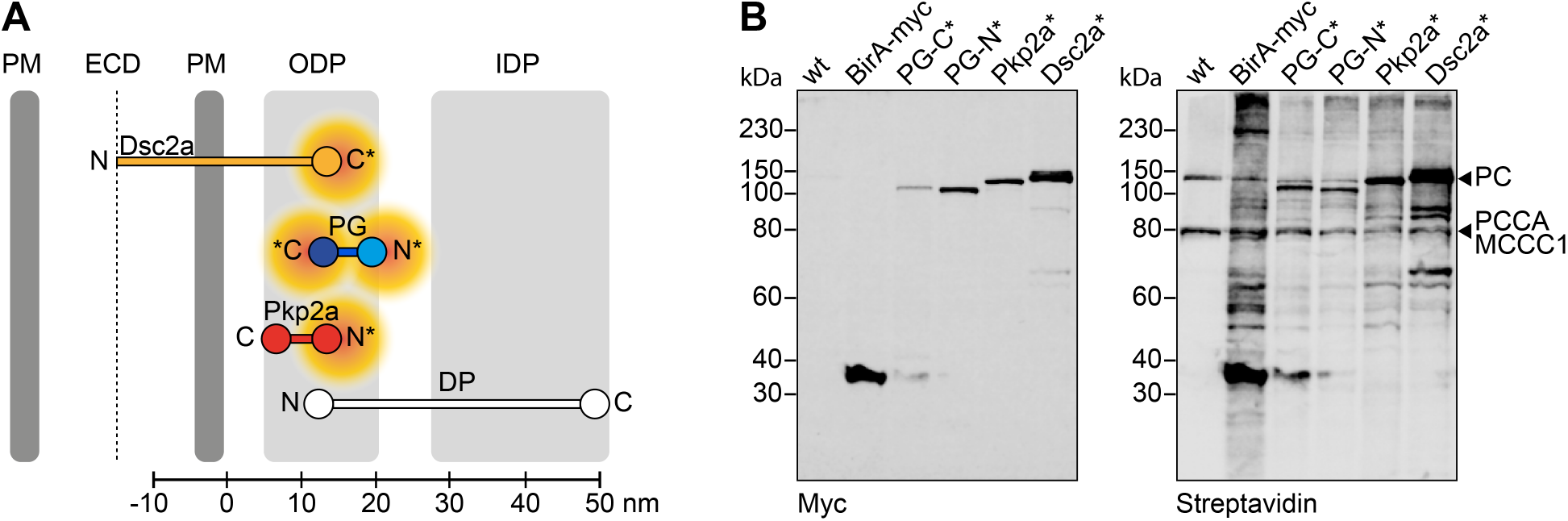
BioID constructs to generate desmosome proximitome. **(A)** Schematic representation of the biotin ligase (BirA) tagged desmosomal constructs, desmocollin 2a (Dsc2a, yellow), plakoglobin (PG, blue) and plakophilin 2a (Pkp2a, red) within the desmosomal molecular map based on work from North et al. (1999). The asterisk indicates the terminus, which was tagged with BirA-myc (C-terminus) or myc-BirA (N-terminus). The orange cloud indicates the estimated radius of biotinylation by the biotin ligase. DP, desmoplakin; PM, plasma membrane; ECD, extracellular core domain; ODP, outer dense plaque; IDP, inner dense plaque. **(B)** Western blot of MDCK cells stably expressing BirA-myc as control, PG-BirA-myc (PG-C*), myc-BirA-PG (PG-N*), myc-BirA-Pkp2a (Pkp2a*) or Dsc2a-BirA-myc (Dsc2a*) and parental wild-type (wt) cells as a negative control. The cells were cultured confluent for 24 h including 16 h with 100 μM biotin before lysis and separation of biotinylated proteins with streptavidin beads. The blot was probed with a mouse-α-myc antibody to detect expression constructs and fluorescently conjugated streptavidin for biotinylated proteins. Note the bands in wt cells correspond with naturally biotinylated mitochondrial carboxylases, including pyruvate carboxylase (PC), methylcrotonoyl-CoA carboxylase subunit alpha (MCCC1) and propionyl-CoA carboxylase alpha chain (PCCA). Blot is representative of three biological replicates.

Western blotting showed expression of all constructs with expected mobilities: 36.5 kDa for BirA-myc, 118.5 kDa for PG-N* and PG-C*, 134 kDa for Pkp2a* and 129.5 kDa for Dsc2a-BirA-myc (Fig. 1 B, left blot). Incubation with fluorescently conjugated streptavidin confirmed the biotinylation capacities of the constructs (Fig. 1 B, right blot) with the most intense bands representing the BioID baits and other bands biotinylated preys. Naturally biotinylated mitochondrial carboxylases and ribosomal proteins (Fig. 1B, table S2 and S3), which can be biotinylated during protein translation, were excluded from further MS analyses. All baits colocalised with DP-positive structures (Fig. S1, left panel), showing that the BirA* tag did not hinder subcellular targeting to desmosomes. Staining for biotinylated proteins revealed distinct distributions in close proximity of the bait proteins, except for a stronger nuclear signal in the BirA-myc-expressing cells (Fig. S1, right panel). These data demonstrated that BirA*-tagged desmosomal proteins localised at the expected cell-cell junction sites where they biotinylate their neighbourhood.

When desmosomes first form between MDCK cells they are Ca^2+^-dependent but the majority become hyper-adhesive within 5 days of confluent culture (Wallis et al., 2000). To determine proximal interactors of each desmosomal BioID bait depending on the desmosomal adhesion state, MDCK cells stably expressing BioID constructs were cultured confluent for 1 day (Ca^2+^-dependent condition) or 5 days (hyperadhesive condition). The proximal proteins (preys) were identified using label-free quantitative mass spectrometry and analysis of the raw data was performed using ion intensity-based quantification with MaxQuant against the Canis lupus familiaris proteome. We used SAINTexpress to identify high-confidence bait-prey proximal interactions with BirA-myc as a negative control. To predict “true” proximity interactors, a stringent Bayesian false discovery rate (BFDR) of ≤ 0.05 was employed and the respective human orthologs were obtained for subsequent analyses. This resulted in a data set with a total of 189 proximal proteins, which included a combination of canonical desmosomal proteins, desmosome-associated proteins and proximal proteins of the BioID baits in ostensibly more distal subcellular locations (Fig. 2 A and tableS1).

**Fig. 2.**
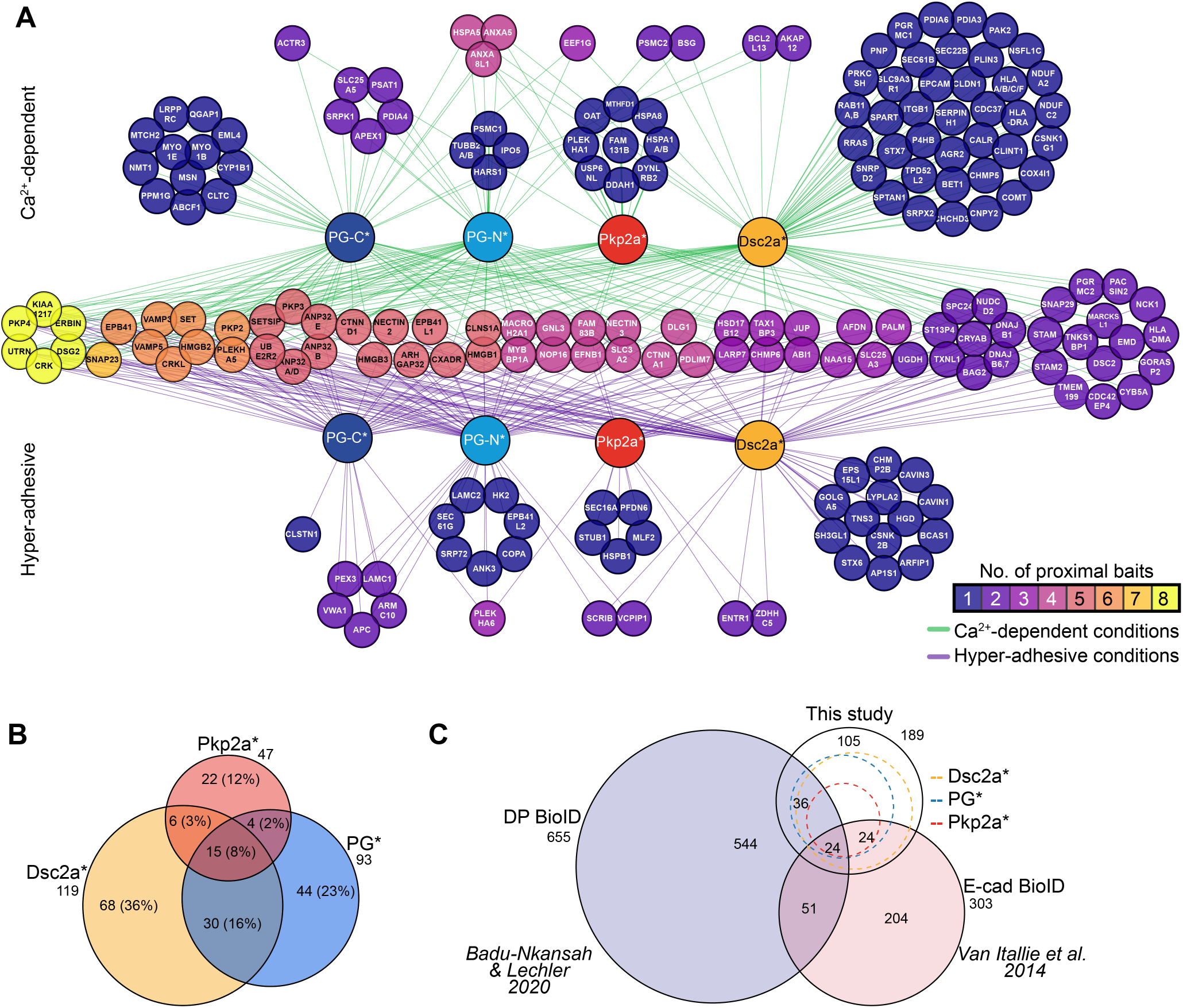
Overview and gene ontology analysis of the partial proximitome of Ca^2+^-dependent and hyper-adhesive desmosomes. **(A)** Network of proximal proteins (gene names of human orthologs) to BirA-myc tagged desmosomal constructs, plakoglobin-BirA-myc (PG-C*, dark blue), myc-BirA-plakoglobin (PG-N*, light blue), myc-BirA-plakophilin 2a (Pkp2a*, red) and desmocollin 2a (Dsc2a*, yellow). Proximal proteins on the top are those identified in samples of MDCK cells cultured confluently for 1 day and thus with Ca^2+^-dependent desmosomes (green edges), those on the bottom were found in samples of cells cultured confluently for 5 days and thus hyper-adhesive desmosomes (magenta edges) and those in the middle were detected in both samples, all with a high-confidence cut-off of BFDR ≤ 0.05Proximal prey proteins are presented using a colour-coded scale indicating the number of proximal baits. **(B)** Area-proportional Venn diagram of proximal proteins to Dsc2a*, Pkp2a* and PG* (combined PG-C* and PG-N*). **(C)** Area-proportional Venn diagram comparing the prey proteins of Dsc2a*, PG* and Pkp2a* to those found in proximity to desmoplakin (DP; Badu-Nkansah et al., 2021) and the adherens junction receptor E-cadherin (E-cad; Van Itallie et al., 2014). Number of proximal proteins are indicated.

We integrated the 8 individual data sets of Dsc2a*, PG-N*, PG-C* and Pkp2a* from both Ca^2+^-dependent and hyperadhesive conditions into a single network to generate a desmosome proximitome consisting of a total of 259 proteinprotein interactions (Fig. 2 A). The overlap between the prox-imitomes of the combined data for the various baits is shown in Fig. 2 B. The network illustrates both the distinct and common proximal proteins to each bait in relation to the desmosomal adhesion state and is organised as follows. The preys in the horizontal middle of the scheme associated with the baits in both desmosomal adhesion states. These “constant” proteins represent 39% (74 proteins) of the total. The colour index indicates how many of the baits shared the preys, with the proteins on the left (yellow) shared by all baits and those on the right shared only one of the baits. The proteins on the upper side of the schematic (78 or 41%) were found only in proximity during Ca^2+^-dependence, whereas those at the bottom (37 or 20%) were uniquely associated during hyperadhesion. The large pools of proteins uniquely associated with particular adhesion states reveal a dramatic change in the neighbourhood of the desmosomal proteins as well as a substantial reduction in their number during desmosome maturation (152 proteins enriched in Ca^2+^-dependent samples to 111 proteins in hyper-adhesive samples).

When comparing the data sets for the different baits, more than half of the proximal proteins (71%, 134 proteins) were unique to a specific bait, whereas 55 proteins (29%) were identified by at least two different desmosomal baits and may therefore represent shared pivotal desmosomal components (Fig. 2 B and table S1).

Comparison with the recently published desmoplakin (DP) proximitome in mouse keratinocytes revealed that 32% (60 proteins) of proximal proteins were shared with our Dsc2a, PG and Pkp2a bait data sets (Fig. 2C and Fig. S2A). These included known desmosomal components, such as Pkp3 and Dsg2, as well as other reported interactors such as erbin and Crk family of adaptor proteins (Badu-Nkansah and Lechler, 2020; Harmon et al., 2013). Among the interactors in the DP proximitome were keratins, intermediate filament proteins that were not significantly enriched in our data sets in relation to the BirA-myc control, reflecting the specific function of DP in connecting intermediate filaments to desmosomes. Further comparison with BioID data comprising proximal proteins of the adherens junction protein E-cadherin in MDCK cells (Van Itallie et al., 2014) revealed 48 mutual proteins (25%) with our data sets (Fig. 2 C and Fig. S2A). The majority of proteins shared with E-cadherin were identified in association with Dsc2a* (39/48) and included trafficking proteins such as syntaxin 7 (STX7) and sec61 translocon β (SEC61B), which may suggest that the both adhesion receptors follow similar routes during the assembly of cell adhesion junctions.

### Functional enrichment analysis of biotinylated proteins

To gain unbiased insight into the diverse functions of the proximity interactors, over-representation analysis of the combined proximitomes of both adhesion states was carried out (Fig. 3 and S2 B, table S4). This allowed investigation into whether the BioID data sets associated with particular cellular components, molecular functions or broader biological functions as represented by current gene ontology (GO) annotations are statistically over-represented among the proximal proteins. A striking observation to emerge from all three analyses was the great similarity of the proximitomes of Dsc2a* and PG* contrasting with the substantially distinct Pkp2a* proximitome.

**Fig. 3.**
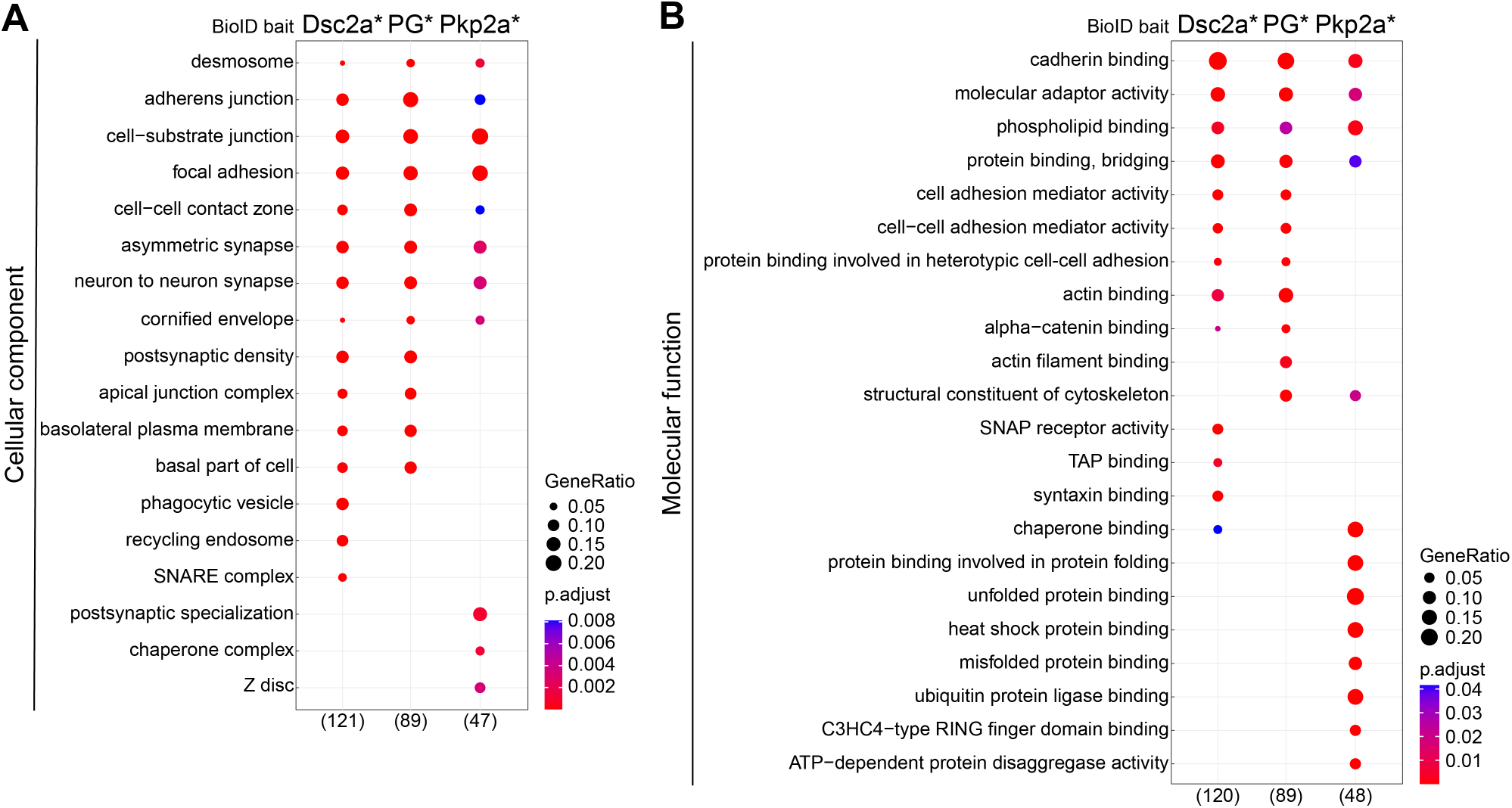
Ontological analysis of desmosomal proximitomes. Functional enrichment analysis of proteins identified by BirA-myc tagged desmocollin 2a (Dsc2a*), plakoglobin (PG*; merged data plakoglobin-BirA-myc and myc-BirA-plakoglobin data) or plakophilin 2a (Pkp2a*) BFDR ≤ 0.05 The top ten overrepresented GO terms under cellular component **(A)** or molecular function **(B)** category are shown. The number of annotated proteins is shown in brackets. (Note 7 protein hits encompassed proteins groups of which all members were included for GO analysis because we could not distinguish between different isoforms. For details see Materials and Methods.) p.adjust, adjusted P value. GeneRatio, proportion of total proteins identified in each GO term.

Cellular component analysis revealed enrichment of terms related to cell-cell junctions, including “desmosome”, “adherens junctions” and “apical junction complex” (Fig. 3 A). Cell-extracellular matrix (ECM) adhesion terms were also identified (“focal adhesion” and “cell-substrate junction”), possibly reflecting miscellaneous targeting of baits and/or shared components and regulatory pathways as reported for Pkp2, integrin β1 (ITGB1) and p120-catenin (CTNND1; Koetsier et al., 2014; Yamamoto et al., 2015).

Similarly, analysis of molecular functions demonstrated some common terms between the data sets of all baits included “cadherin binding”, “molecular adaptor activity”, “protein binding, bridging” and “phospholipid binding”, which correspond with desmosomes being multi-protein complexes associated with the phospholipid plasma membrane (Fig. 3 B). Among lesser-annotated molecular functions was “four-way junction DNA binding” which included high mobility group box proteins (HMGBs) which are chromosomal proteins involved in transcription, replication, recombination and DNA repair (Hock et al., 2007). The majority of molecular function terms for Pkp2a* were distinct from those of Dsc2a* and PG*, suggesting a separate signalling role for Pkp2a. Most of the preys based on GO annotations associated with the appropriate folding of proteins include chaperones and heat shock proteins that are further analysed and discussed below.

In conclusion, the two types of dynamic behaviour of desmosomal components that we recently demonstrated (Fülle et al., 2021) is reflected in the divergent nature of their proximitomes, with those of the two stable components Dsc2a and PG being similar in their molecular neighbourhood while that of the much more dynamic Pkp2a is markedly distinct.

### Maturation analysis outlines neighbourhood differences between desmosomal components and shows substantial changes as desmosomes become hyper-adhesive

To gain deeper insight into the possible functional significance of the proximitome during the switch in adhesion states, we performed two-dimensional hierarchical clustering and GO enrichment analyses of the clusters of prey proteins (Fig. 4 A, B; Fig. S3 A). Such hierarchical clustering helped us to separate the data into clusters of similarity related to the adhesion state of the desmosomes. This resulted in eleven clusters, which fell into three categories with respect to desmosome maturation from Ca^2+^-dependence to hyper-adhesion: those that are present throughout, the “constant” proteins (cluster 7, 9 and 10); those that are present exclusively during the Ca^2+^-dependent phase (clusters 3, 5, 11 and a slight bias in cluster 2), and those that are present exclusively during the hyper-adhesive phase (clusters 1,4, 6, and 8). Moreover, except for cluster 10, the clusters mostly focussed around specific desmosomal baits: clusters 3, 4, 6, 7 and 11 were focussed mainly on PG*, clusters 5, 8, and 9 mainly on Dsc2a*, and clusters 1 and 2 mainly on Pkp2a*.

**Fig. 4.**
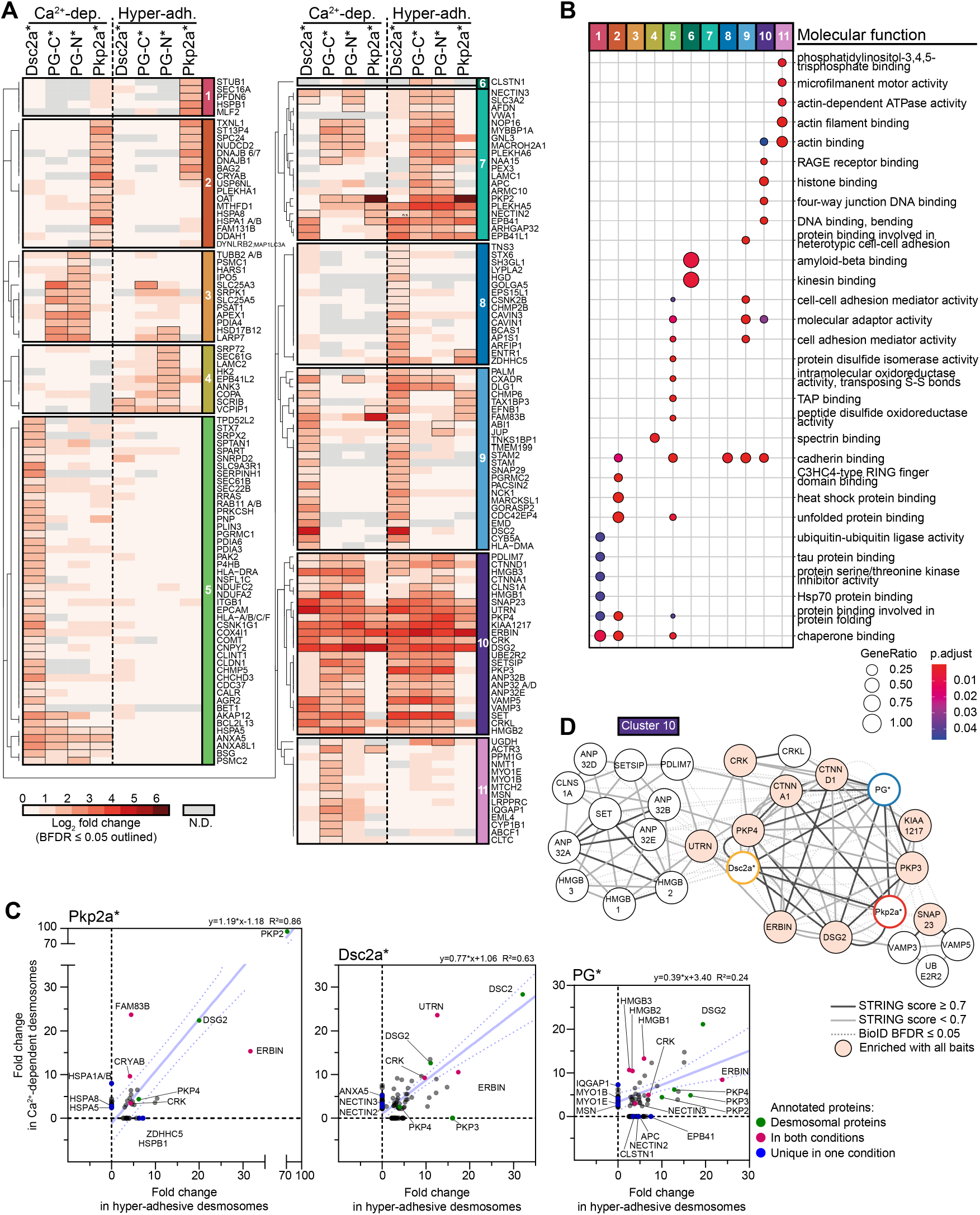
Differences in the proximitome of Ca^2+^-dependent and hyper-adhesive desmosomes revealed by hierarchical clustering. **(A)** Hierarchical clustering was performed on the 189 proteins identified in the partial desmosomal proximitome and displayed as a heatmap. Prey proteins were clustered based on the Jaccard distance of significant (BFDR ≤ 0.05) and non-significant (BFDR > 0.05) interactions across all baits i.e. desmocollin 2a-BirA-myc (Dsc2a*), myc-BirA-plakoglobin (PG-N*), plakoglobin-BirA-myc (PG-C*), myc-BirA-plakophilin 2a (Pkp2a*) and both conditions i.e. 1 or 5 days confluently cultured MDCK cells stably expressing BioID baits thus with either Ca^2+^-dependent (Ca^2+^-dep.) or hyper-adhesive desmosomes (Hyper-adh.) respectively. Log2 fold-change enrichment over BirA-myc control of prey proteins are displayed as a heatmap. Dendrograms were split to identify clusters of preys (1—11). N.D. = not detected. **(B)** GO enrichment analysis of the clusters in A. The top 5 overrepresented terms under the molecular function category per cluster are shown. **(C)** Network analysis of bait-prey interactions of cluster 10 in A. Nodes of bait proteins are colour codes with Pkp2a* in red, Dsc2a* in yellow and PG* (merged PG-N* and PG-C*) in blue. Nodes of prey proteins are shaded in light red when they were present in all bait data sets. Edges indicate protein-protein interactions: solid dark grey lines indicate a STRING score ≥ 0.07 solid light grey lines a STRING score below 0.7 and dotted red lines BioID proximity with a BDFR ≤ 0.05 presented in this study. **(D)** Scatter analysis highlighting changes of prey proteins during desmosome maturation. Scatter plots showing the relationship of of prey proteins in cells with either Ca^2+^-dependent or hyper-adhesive desmosomes with either Dsc2a*, Pkp2a* or PG* as a bait (SAINT fold change enrichment over BirA* control). The 95% confidence interval of the linear regression line is displayed as dotted confidence band.

To determine how the proximal proteins to PG* were shared between its N- and C-termini, we also generated a heatmap following hierarchical clustering of the data sets at both adhesion states for PG-N* and PG-C* (Fig. S3 B and C). 39 (42%) of the total 93 proteins were detected by both PG-N* and PG-C* irrespective of the desmosomal adhesion state (Fig. S2 C). This substantial overlap may be because the range of biotinylation of BirA* probably covers the central armadillo (arm) domain of PG, which binds both DCs (arm 1-3), Pkps and DP and other shared proximal interactors (Al-Jassar et al., 2013).

Whilst some proteins were detected irrespective of the desmosomal adhesion state and thus could be classified as “constant”, it is clear from the heatmaps that many of these exhibited stoichiometric changes during the maturation process (Fig. 4A and Fig. S3 B). To examine this possibility in more detail we generated scatter plots that show the quantitative relationships of prey proteins depending on the adhesion state (Fig 4 C and Fig. S3 C). These analyses indicate that while some properties of desmosomes persist in all states, others change substantially with the adhesion status of the junction.

To gain further information about how our experimental data overlay with existing databases we added STRING based interaction networks to each cluster (Fig. 4 C and Fig. S4; Szklarczyk et al., 2021). We have selected some of these networks for further discussion below; others will serve as undiscussed resources awaiting future analysis.

### Constant proteins: some novel interactions and functional insights

Proteins that are present throughout the maturation process are likely to be continuously involved in desmosome function. Cluster 10 contains proteins that were observed proximal to all baits (Fig. 4 C) with the fewest changes between desmosomal adhesion states; cluster 9 proteins were mostly continuously present close to Dsc2a*, and cluster 7 proteins were enriched proximal to PG* in both conditions (Fig. S4). These clusters comprised canonical desmosomal proteins (PKP3, PKP4 and DSG2; cluster 10), proteins of other cellcell adhesion complexes such as the nectin 2 and 3 (cluster 7) and *α*- and p120-catenin (CTNNA1 and CTNND1; cluster 10) of adherens junctions. CRK and CRKL also contributed to an enrichment of cell-cell adhesion-relevant GO annotations. Both Crk and CrkL were shown to localise to desmosomes and to be critical for desmosome integrity, in particular for the localisation of PG to the desmosomal plaque in the epidermis of Crk/CrkL knock-out mice (Badu-Nkansah and Lechler, 2020). These proteins were over-represented in cell-cell adhesion-relevant terms “cadherin binding” and “molecular adaptor activity” and outline the proximity with potential signalling crosstalk to the neighbouring cadherin- and nectin-associated adherens junctions (Barron et al., 2008; Lewis et al., 1997; Vasioukhin et al., 2000; Yoshida et al., 2010). A novel association appeared in Cluster 9 that hinted at some crosstalk of desmosomes with tight junctions. This cluster depicted mainly Dsc2a*-associated constant proteins and included CXADR (Coxsackie virus and adenovirus receptor (CAR)) that is necessary for tight junction integrity and trans-epithelial leukocyte migration (Fig. 4 A and Fig. S4; Cohen et al., 2001; Morton and Parsons, 2011). Cluster 9 also included NCK adaptor protein 1 (NCK1), which is involved in signal transduction from receptor tyrosine kinases and the actin cytoskeleton (Buday et al., 2002; Li et al., 2001).

It should be noted that, unlike desmosomes, the majority of these cell-cell adhesion proteins are linked to the actin cytoskeleton. In contrast to the intermediate filaments that are associated with desmosomes, the actin cytoskeleton contains motor proteins that mediate contractility. The proximity of the two systems may be consistent with some of the actin dependent mechanisms that regulate desmosome function, including the actin dependent internalisation of desmosomes (Fülle et al., 2021; McHarg et al., 2014).

A group of novel proteins in cluster 10 that remained in the proximitome of desmosomes irrespective of the adhesion state belonged to terms associated with DNA and histone binding (“histone binding”, “four-way junction DNA binding” and “DNA binding, bending”) (Fig. 4 B), including the high mobility group proteins 1-3 (HMGB 1-3) and the acidic nuclear phosphoprotein 32 family members A, B, D and E (ANP32 A/B/D/E). Network analysis of cluster 10 including computational STRING-based evidence scores revealed the strong relation of adhesion-relevant proteins with desmosomal components and the lesser-known relationship of this subnetwork to the nuclear proteins (Fig. 4 C). These data suggest a potentially important yet rather unexplored functional role of desmosomes in the regulation of transcription factors. HMGB1/2/3 are drivers of various cancers and have nuclear chromatin binding functions, regulation of both telomerase activity and transcription, as well as cytoplasmic and cytokine-like functions, driving inflammation and proliferation when released from cells (Sharma et al., 2008; reviewed by Wen et al., 2021). Furthermore, HMGB proteins were shown to be regulated by both PG and β-catenin (β-catenin/WNT pathway) (reviewed by Wen et al., 2021). Additionally, considering their cytokine-like functions, they may play a pivotal role in regulating cell proliferation in relation to IF-associated cell junctions. It is interesting to speculate that PG is also directly involved in the regulation of HMGB1/2/3 providing a reciprocal signal of the adhesion state of desmosomes to the nucleus. Alternatively, PG might compete with β-catenin in the regulation of HMGBs. It should be noted that, whilst the HMBGs are “constant” proteins, their prey-to-bait ratios are substantially greater during the Ca^2+^-dependent phase than during hyper-adhesion (Fig 4 D) so it may be that their functional significance is greater during keratinocyte activation (e.g. wound healing (Tamai et al., 2011)) and declines as desmosomes mature.

In contrast, erbin is a constant protein that shows an increase prey/bait ratio in hyper-adhesion. Intriguingly, erbin was the most enriched protein within the PG* data sets during hyperadhesion and was also increasingly enriched in proximity to Dsc2a* and Pkp2a* (Fig. 4 A, D: Fig. S3 C). Erbin is an adaptor protein and a critical regulator of the localisation and signalling of the receptor tyrosine-protein kinase erbB-2 (ERBB2 /HER2) and Ras-mediated MAPKs signalling pathways (Borg et al., 2000). It has also been associated with numerous other signalling pathways and several binding partners have been identified including the desmosomal accessory protein Pkp4 (also known as p0071) (reviewed by Dan et al., 2010). Interestingly Pkp2,3 and 4 follow a similar increased presence in the proximitome as erbin. Since the interaction of erbin and Dsg1 was shown to disrupt SHOC2-Ras-Raf complexes (Harmon et al., 2013), the neighbourhood could play critical role in signalling cell quiescence as cell density increases and desmosomes progressively switch to a hyper-adhesive state.

A striking discovery was the finding of multiple heat shock proteins (HSPs) and other chaperones in proximity to Pkp2a*, principally as constant proteins and additional ones during Ca^2+^-dependence (cluster 2 in Fig. 4 and Fig. S4). The HSP family encompasses several sub-groups of chaperones, which are particularly important in response to cellular stressors (Kampinga et al., 2009). The individual roles of HSPs are manifold, but their main function involves folding and stabilising newly formed proteins. The novel finding suggests a potential connection between HSP and desmosomes and encourages further investigation.

### Ca^2+^-dependence: Possible roles in desmosome assembly

The Ca^2+^-dependent phase is associated with desmosome assembly, a process which appears to continue for at least 36 hours subsequent to its initiation in MDCK cells (Mattey et al., 1990). Desmosomes also become Ca^2+^-dependent upon epithelial wounding (Kimura et al., 2007) and during wound healing, in areas where keratinocytes become activated for migration that requires motility and desmosome turnover. At this time desmosomes are less stable than during hyperadhesion (Bartle et al., 2020; Fülle et al., 2021; Kimura et al., 2007; Wallis et al., 2000) and both assembly and internalisation require crosstalk with the actomyosin machinery (Fülle et al., 2021; Lewis et al., 1997; Vasioukhin et al., 2001).

Such crosstalk is reflected particularly in group of prey hits of PG* that were enriched in Ca^2+^-dependent conditions (cluster 11 in Fig. 4 A; Fig. S3 B and C; Fig. S4) and were mostly linked to actin associated terms (“actin-dependent ATPase activity”, “actin filament binding”, “actin binding” and “microfilament motor activity”) (Fig. 4 B; cytoskeleton in Fig. 5). Members of these groups were the actin-based motor proteins myosin 1B and E (MYO1B /E), actin-plasma membrane cross-linker moesin (MSN), Arp2/3 complex subunit actin-related protein 3 (ACTR3 i.e. ARP3) and the IQ motif containing GTPase activating protein 1 (IQGAP1), all of which are known regulators of the actin cytoskeleton (Brandt and Grosse, 2007; Cheng et al., 2012; Goley and Welch, 2006; Niggli and Rossy, 2008; Salas-Cortes et al., 2005).

**Fig. 5.**
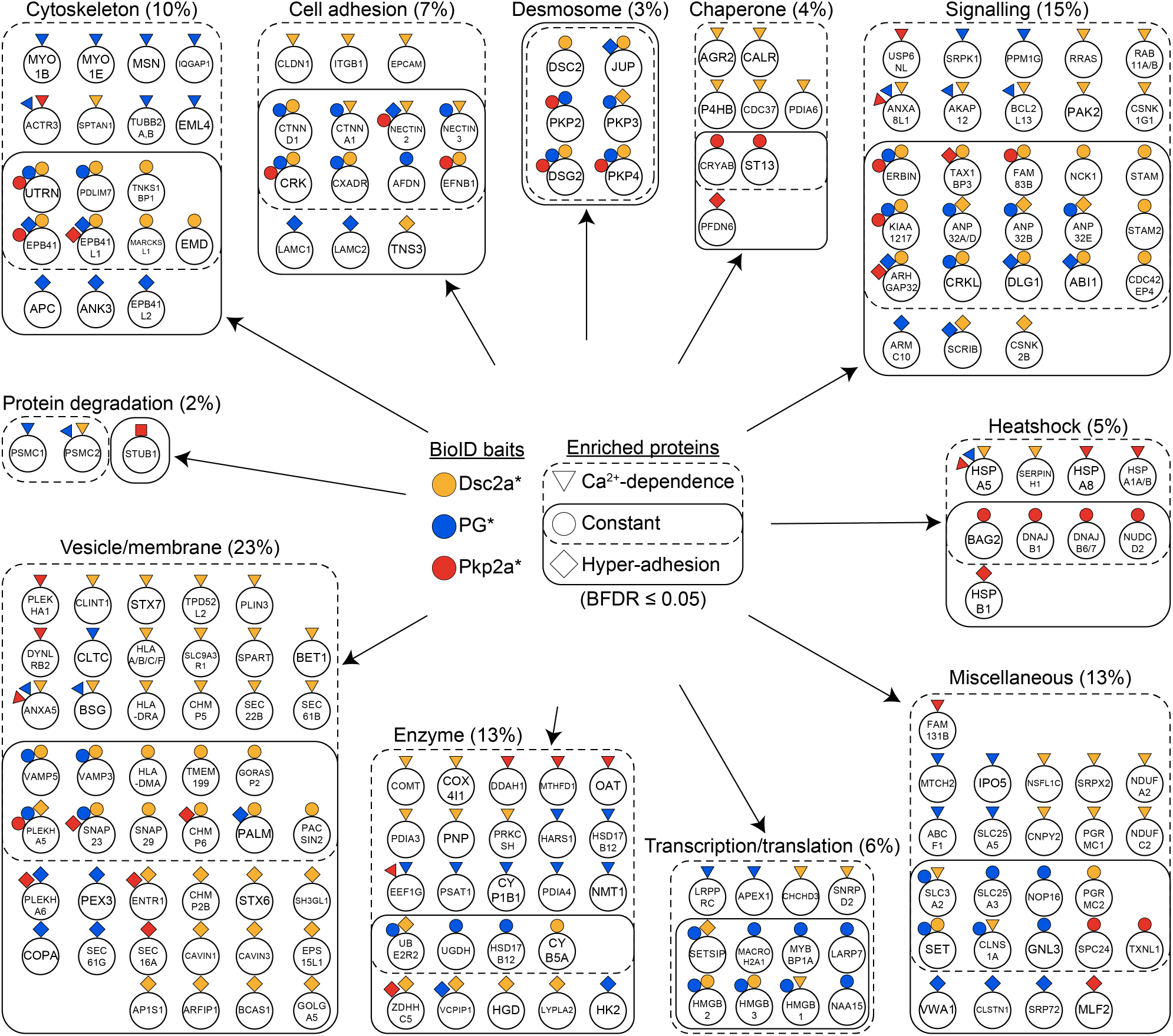
Functional characterisation of desmosome proximal proteins. The function of the 189 proximal proteins (BDRF ≤ 0.05) of BirA-myc tagged desmocollin 2a (Dsc2a*), plakoglobin (PG*; merged data plakoglobin-BirA-myc and myc-BirA-plakoglobin data) or plakophilin 2a (Pkp2a*) were annotated based on primary literature and the Human Protein Atlas and classified into the indicated categories. The proximal baits are represented on each prey with the correspondingly coloured geometric shapes (Dsc2a* in yellow, PG* in blue and Pkp2a* in red). Boundaries and geometric shapes indicate whether the prey proteins were significantly enriched in data curated from MDCK cells cultured confluently for either 1 day and thus with Ca^2+^-dependent desmosomes (dotted line, triangular bait), for 5 days and thus hyper-adhesive desmosomes (solid line, square bait) or enriched in both culture conditions (overlap, circular bait). The percentages of the total protein number are indicated in brackets

Another large fraction of Dsc2a* proximal interactors (cluster 5 in Fig. 4 A and Fig. S4) fall into similar categories as actin regulatory proteins and contain the Rho GTPases and related proteins, such as Cdc42 effector protein 4 (CDC42EP4), RAS related (RRAS) and p21 (RAC1) activated kinase 2 (PAK2). All these prominent actin modulators are associated with membranes and possibly have a role in the concomitant assembly of desmosomes and adherens junctions (Etienne-Manneville and Hall, 2002). In line with the notion that desmosomes increasingly separate from other junctions (see above) during maturation is the finding that other junctional receptors such as integrins (INGB1), claudins (CLDN1) and EpCAM (EPCAM) are exclusively present in the proximitome of Dsc2a under Ca^2+^-dependent conditions.

A smaller fraction of proteins in Ca^2+^-dependent desmosomes was uniquely linked to Pkp2a (cluster 2 in Fig. 4 A; Fig. S4), most of which belong to heatshock proteins (compare with Fig. 5). Amongst them were family members of Hsp70 proteins (HSPA1A, HASPA1B, HSPA5, and HSPA8). Gao and Newton showed that Hsp70 is involved in the stabilisation and re-phosphorylation of PKCs (Gao and Newton, 2002). These results are consistent with the importance of PKCs in the regulation of the desmosomal adhesion state (Bass-Zubek et al., 2008; Kimura et al., 2007; Kröger et al., 2013; Thomason et al., 2012; Wallis et al., 2000). Translocation of PKC*α* to desmosomes and phosphorylation of DP was shown to precede Ca^2+^-dependence of desmosomes (Kröger et al., 2013; Wallis et al., 2000). Furthermore, PKC-scaffolding annexins were present particularly under Ca^2+^-dependent conditions for all baits (cluster 5 in Fig 4A: Fig. S4). Annexins are known substrates and scaffolding proteins of multiple PKC isoforms (Hoque et al., 2014), so their juxtaposition to desmosomes (ANXA5, vesicle/membrane fraction, Fig. 5) might contribute to the PKC*α*-mediated adhesion switch of desmosomes shown by Wallis et al. (2000).

One of the largest fractions uniquely associated with different adhesion states is linked to vesicles and membranes, predominantly in the neighbourhood of the transmembrane protein Dsc2a (clusters 5 and 8 in Fig. 4 A; Fig. S4; Fig. 5). Many of these proteins are involved in membrane organisation, transport of vesicles or membrane channels. How the dramatic switch in membrane neighbourhood contributes to desmosome function or, vice versa, how desmosomes may contribute to the changes of this sub-cellular compartment, with potentially strong impact cellular function, remains to be elucidated.

### Hyper-adhesion: Functions of mature desmosomes

Hyper-adhesion is the condition of most desmosomes in tissues and thus the interacting proteins found here most probably represent the stable state of desmosomal function. The current analysis reveals several novel interactions of desmosomes as well as some confirmatory indications.

Cluster 8 included Dsc2a* preys uniquely enriched under hyper-adhesive conditions, many of which are linked to the localisation of membrane associated proteins (Fig. 4 A and Fig. S4). Those included proteins involved in caveolar endocytosis including caveolae associated proteins 1 and 3 (CAVIN1/3) and epidermal growth factor receptor pathway substrate 15 like 1 (EPS15L1) (Kovtun et al., 2015; McMahon et al., 2009). These results are consistent with results from Brennan et al. (2012) showing co-localisation of Dsg2 with Cavin1. In their study, authors proposed a caveolae-dependent internalisation and subsequent degradation of proteolytically truncated Dsg2, critical for the maintenance and stability of desmosomes. Our results suggest that this process is important for matured hyper-adhesive desmosomes.

Clusters 4 and 6, and to some extent cluster 7, represent particularly PG* preys enriched in hyper-adhesion (Fig. 4 A; Fig. S3; Fig S4). Calsyntenin-1 (CLSTN1, alone forming cluster 6, was discovered in postsynaptic membranes (Vogt et al., 2001) and belongs to the superfamily of cadherins, thus potentially containing a PG binding site. The tumour suppressor adenomatous polyposis coli protein (APC) appears in the vicinity of PG (cluster 7; Fig. S4). APC was previously found to bind PG, and together with the scaffolding protein axin, it is possibly involved in the degradation and thus the homeostasis of PG levels of mature desmosomes (Kodama et al., 1999). Cluster 4 (Fig. S4) included the scaffolding proteins ankyrin 3 (ANK3 also known as Ankyrin G) and erythrocyte membrane protein band 4.1 like 2 (EPB41L2). These are part of the cortical spectrin-actin networks as they are both able to bind spectrin (Baines et al., 2014). The importance of an intact cortical actin network for the turnover and stability of the desmosomal complex was recently shown by knockout and knockdown of α-adducin, a component of the spectrin-actin network, in keratinocytes (Hiermaier et al., 2021; Rötzer et al., 2014). Our results appear consistent with the view that the development of desmosomal hyperadhesion is part of a process of stabilisation of the cortical cytoskeleton and the quiescence of cells, also involving spectrin and associated proteins (Ghisleni et al., 2020; Patel et al., 2019).

### Re-categorisation of protein function

GO analysis allows rapid insight into potential cellular and molecular functions of preys based on a vast array of experimental data. However, since GO annotations continuously evolve as experimental data become available (Gaudet and Dessimoz, 2017; Tomczak et al., 2018), GO enrichment analysis can be affected by annotation bias (annotations from only few well-studied genes) or literature bias (few articles that contribute disproportionally to annotations). Such bias may explain the relatively high number of neurology related GO terms even though our data relate to epithelial cells. Examples of such terms include “postsynaptic density” and “asymmetric synapse” (Fig. 3 A). Proteins with these annotations include p120 catenin, Nectin 3 and the Rho GTPase-activating protein 32 (ARHGAP32 i.e. RICS). The latter is involved in the β-catenin-N-cadherin receptor signalling and critical for synaptic adhesions of neurons (Okabe et al., 2003).

To minimise such bias we complemented the GO analysis with our own map of manually annotated protein functional terms based on “The Human Protein Atlas”, an omic database that provides a wider range of information. Ten functional categories, which encompassed all preys, evolved during the process of manual annotation: cytoskeleton, cell adhesion, desmosome, chaperone, signalling, heat shock, transcription/translation, enzyme, vesicle/membrane, protein degradation, and miscellaneous when multiple categories applied or the protein function was unknown (Fig. 5). Whilst this complementary functional classification does not comprise a statistical “enrichment analysis” and could be extended in further subgroups, it highlights functional terms that may be more comprehensive in relation to epithelial cell functions. Figure 5 also clearly illustrates the changes in the environments of these major desmosomal proteins during the process of maturation form Ca^2+^-dependence to hyper-adhesion.

### Conclusion

The picture that emerges is of a junction which, though fundamentally stable and mechanical in function, possesses extensive signalling capacity and which changes its interactome considerably according to whether it is in a less stable, immature state or a more stable, more strongly adhesive mature state. Furthermore, our data support our recently proposed concept of desmosomal dualism as the characteristics observed here revealed of Pkp2a, the constantly dynamic component, differ from those of Dsc2a and PG, which are much more stable (Fülle et al., 2021). In figure 6, we summarise this and emphasise some of the major points that emerge from our results.

**Fig. 6.**
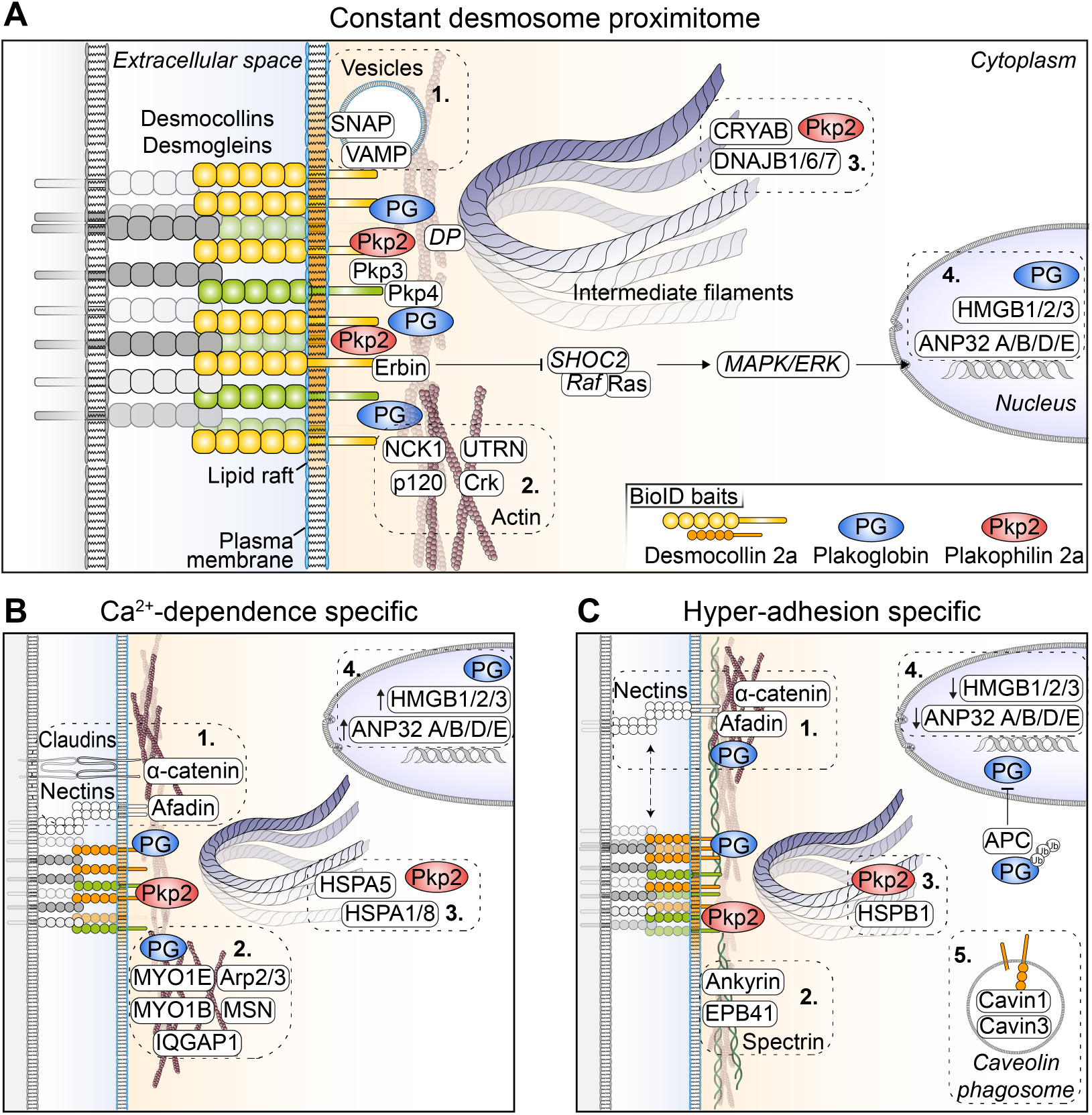
Schematic diagrams of the desmosome proximitome and differences between the Ca2+-dependent and hyper-adhesive phases. **(A)** Constant proteins that are present in both phases, **(B)** proteins present exclusively during Ca^2+^-dependence, **(C)** proteins present exclusively during hyper-adhesion. Not all significantly enriched proximal proteins are shown. Those written in italics were not significantly enriched but have been inserted for discussion purposes. For further details see text where the numbered boxes are referred to. (Small solid arrows indicate increased or decreased protein ratios.)

During junction assembly, desmosomal proteins are closely associated with those of adherens junctions (AJ) but subsequently they segregate into separate junctions. This compartmentalisation of junctions is clearly indicated in our findings where the AJ proteins nectin 2 and 3 and α-catenin are close to Dsc2a when desmosomes are Ca^2+^-dependent (Box 1. in Fig. 6 B), but become spatially more distinct as desmosomes mature to hyper-adhesion (Box 1. in Fig. 6 C). These proteins, however, retain proximity to PG, possibly reflecting the capacity of PG to locate at adherens junctions as well as desmosomes (Lewis et al., 1997). This highlights the fact that both PG and Pkp2 can reside in non-desmosomal locations.

Spatial separation of junctions is accompanied in desmosomes by intriguing changes in the associated actin binding proteins. Different actin-binding proteins interact, especially with PG, depending on the adhesion state (Box 2 in Fig 6 B; Boxes 1 and 2 in Fig. 6 C). These include adaptor proteins (e.g. p120 catenin, α-catenin and afadin), and those involved in endocytosis (e.g. myosin 1E [MYO1E]) or in actin dynamics (e.g. NCK1 and Arp2/3). A shift towards cortical spectrin associated proteins (e.g. ankyrin and EPB41) suggests a remodelling and stabilisation of the cortical spectrin-actin network concomitant with the maturation of desmosomes (2. In C). We also noted, as expected, the lack of interaction of the Dsc2a, PG and Pkp2a baits with keratin proteins. However, the HSPs CRYAB and HSPB1, which associate with Pkp2a as constant and hyper-adhesion proteins respectively, have been shown elsewhere to interact with intermediate filaments, possibly preventing their aggregation (Kumarapeli and Wang, 2004; Perng et al., 1999). Also noteworthy is that these two Pkp2-associated HSPs change as desmosomes mature (Boxes 3 in Fig. 6 B and C). It is interesting to speculate whether this may indicate a Pkp2a-regulated change in the nature of keratin filament interaction with desmosomes, or whether it represents a separate interaction between Pkp2a and keratin filaments in a location remote from desmosomes as shown in Fig. 6.

The desmosomal cadherin desmoglein 1 has been shown to regulate the Ras/MAPK/ERK pathway and thus transcriptional activity in epidermal keratinocytes (Harmon et al, 2013). In view of this we speculate that the ratios of erbin increasing near hyper-adhesive desmosomes may indicate that desmosomes regulate cell quiescence via the same pathway (Fig. 6 A, italics). The association of other nuclear proteins with desmosomes, in ratios decreasing with hyper-adhesion (Box 4 in Fig. 6), suggests further potential for regulation of gene expression by desmosomal proteins. Whether this may involve an extra-desmosomal role for PG or some type of cell junction to nucleus signalling as has been found for tight junction remains to be seen (Balda et al., 2003). Proximity to APC may potentially reduce the cytoplasmic and nuclear levels of PG and thus affect desmosome-related signalling (Kodama et al., 1999; Ozawa et al., 1995; Rubinfeld et al., 1995).

Desmosomes are embedded in lipid rafts, which are microdomains of the plasma membrane enriched in sphingolipids and cholesterol and which are believed to play a role in desmosome assembly, function and disassembly (Resnik et al., 2011; reviewed by Zimmer and Kowalczyk, 2020). In this context it is interesting that several membrane and vesicle associated proteins are constantly in proximity to Dsc2a (Box 1. in Fig. 6 A). In addition the proximity of cavin 1 and 3 to Dsc2a (Box 5 in Fig. 6 C) suggests homeostasis of desmosomal cadherins through caveolae-dependent internalisation is important in hyper-adhesive desmosomes. Desmosomes are remarkably stable structures (Fülle et al., 2021; Windoffer et al., 2002), with FRAP studies indicating no more than a 20% turnover of Dsc2a within 5 minutes. However, the appearance of cavins in our present studies following biotin labelling for 12 hours indicates a possible continuous turnover of Dsc2a throughout the lifetime of the desmosome.

In closing we would like to stress our belief in the importance of maturation studies such as presented here. Though currently limited, the evidence suggests that the hyper-adhesive state acquired by desmosomes during extended confluent tissue culture resembles that acquired by and maintained by desmosomes in tissues (Garrod et al., 2005; Kimura et al., 2012; Thomason et al., 2012; Wallis et al., 2000) and is therefore highly relevant to the function of desmosomes in vivo. One of our major findings has been the substantial differences between the proximitomes of desmosomes in this hyperadhesive state compared to those that are newly formed.

## ACKNOWLEDGEMENTS

We thank the staff of the Biological Mass Spectrometry Core Facility, in particular Stacey Warwood, Emma-Jayne Keevill and David Knight, and Bioimaging Core Facility, in particular Peter March and Roger Meadows, of the Faculty of Biology, Medicine and Health at the University of Manchester for their support and help with this project. We would extend our gratitude to Jonathan Humphries and Megan Chastney for their help and advice regarding the MS experiments. The Ballestrem laboratory is part of the Wellcome Trust Centre for Cell-Matrix Research, University of Manchester, supported by core funding from the Wellcome Trust (grant number 203128/Z/16/Z). We further want to acknowledge the Biotechnology and Biological Sciences Research Council (BBSRC; grant number BB/R001707/1, BB/R014361/1) and the Wellcome Trust (grant number 202923/Z/16/Z) for funding this project. The authors declare no competing financial interests.

## Materials and Methods

### Reagents

Primary antibodies used at the indicated dilutions for immunofluorescence were mouse α-desmoplakin I and II (clone 11-5F, custom-made by D.R.G. (Parrish et al., 1987)), 1:200; rabbit α-desmoplakin (A303-356A, Bethyl Laboratories), 1:200; and mouse α-myc (clone 9B11, Cell Signalling Technology). Secondary antibodies conjugated to Alexa Fluor 488 or 647 were from Thermofisher (used at 1:500). Alexa Fluor 488-conjugated phalloidin and Alexa 680-conjugated streptavidin were from Life Technologies (both used at 1:500). DAPI readymade solution (Sigma) was used at a concentration of 1 μg/ml. Y-27632 dihydrochloride (Tocris Bioscience) was dissolved in water and used at a final concentration of 50 μM.

### Cell culture

Madin-Darby canine kidney II cells (MDCK; ECACC) (Madin and Darby, 1958) were cultured at 37°C in 5% humidified CO_2_ in high glucose Dulbecco’s Modified Eagle Medium (DMEM; Sigma), supplemented with 10% (v/v) foetal calf serum (FCS; Gibco) and 100 U/ml penicillin and 100 μg/ml streptomycin (P/S; Gibco).

### Cloning

To generate constructs containing BirA-myc, cloning was performed using Gibson Assembly Cloning Kit EE5510S, New England Biolabs (NEB) according to the manufacturer’s protocol. Vectors were linearized with single restriction endonucleases obtained by NEB, and fragments were generated by polymerase chain reaction (PCR) using Phusion High-Fidelity Polymerase (M0530L, NEB) using 35 cycles and 60°C annealing temperature. All primers were designed using SnapGene (GSL Biotech LLC, Chicago, IL) and were synthesised by Eurofins Genomics (Germany) (Table S5). BioID vectors pCDH-BirA-myc and pCDNA3.1-myc-BioID (Addgene no. 35700 (Roux et al., 2012)) were gifts from A. Gilmore (University of Manchester, UK) and C. van Itallie (NIH, US). To generate puro-BirA-myc containing expression vectors, BirA-myc was obtained by PCR using pCDH-BirA-myc as a template with primes BirA-myc.for and BirA-myc.rev and cloned via NcoI site into a custom-made vector pSF (0G394R1, Oxford Genetics) which was modified to have an EIF1a promoter and a puromycin selection marker. To clone myc-BirA, pCDNA.31-myc-BioID and primers myc-BirA.for and myc-BirA.rev were used and cloned into pSF via NcoI site. Plasmid DNA containing desmosome genes were purchased from Addgene, [Dsc2a no. 32233 (Ishii et al., 2001), PG no. 32228 (Palka and Green, 1997) and Pkp2a no. 32230 (Chen et al., 2002)] and used as template for PCRs to obtain their open reading frames (ORF) (Table S5). To construct puro-Dsc2a-BirA-myc and puro PG BirA myc the primers Dsc2a.for and Dsc2a.rev, PG-BirA.for and PG-BirA.rev were used, respectively, for both constructs, the fragments were cloned into HindIII site. Puro myc BirA PG was cloned into puro-myc-BirA via BbvCI site using primers BirA-PG.for and BirA-PG.rev, and Pkp2a was amplified using Pkp.for and Pkp.rev and cloned into puro myc-BirA via BbvCI site creating puro-myc-BirA-Pkp. All ORF sequences were confirmed by Sanger sequencing performed by Eurofins.

### Generation of stable cell lines

Cells were transfected using Lipofectamine LTX transfection reagent, according to the manufacturer’s instructions (Invitrogen). To generate stable cell lines, transfected MDCK cells were selected using 2 μg/ml puromycin (Thermofisher) in SM with medium changes every 2 days for 10 days.

### Immunofluorescence microscopy

Cells were fixed with 4% (w/v) paraformaldehyde in PBS or with 100% ice cold methanol for α-desmoplakin (clone 11-5F) immunostaining. Antibodies were diluted in 1% BSA and added to the cells for 1 h. Images were acquired on a Delta Vision microscope (Applied Precision) with a 60×/1.42 Plan Apo N (Oil) objective and a Sedat Quad filter set, with images collected using a Retiga R6 (Q-Imaging) camera and processed using the FIJI ImageJ software (version 1.53 g; https://fiji.sc/).

### Proximity biotinylation and affinity purification

To mediate proximity biotinylation, cells expressing BirA-myc constructs were seeded at confluent density (1.35×105 cells/cm2) onto 15-cm plastic cell culture dishes for 8 h for Ca^2+^-dependent desmosome formation or 4 d and 8 h for hyper-adhesive desmosome maturation, and then incubated in medium supplemented with 100 μM biotin (B20656, Life Technologies) for 16 h (i.e. total culture time was 1 d or 5 d).

Biotinylated proteins were affinity purified following a protocol adapted from Chastney et al. (Chastney et al., 2020). Cell lysis was performed in a 4°C cold room. The cells were washed thrice with cold 15 ml PBS (without CaCl2 and MgCl2) and lysed with 1.2 ml lysis buffer (50 mM Tris pH 7.4, 500 mM NaCl, 0.4% [wt/vol] SDS, 5 mM EDTA, 1 mM DTT and 1x cOmplete Protease inhibitor cocktail, Roche) for 20 min on a rocker at 4°C. Cells were scraped, transferred in 5 ml tubes (Eppendorf) and 480 μl Triton-X-100 (2% [vol/vol] final concentration) was added. Samples were further lysed on ice by three sonication steps for 30 sec each using a Vibra-Cell VCX750 (Sonics, US) at 20% power. 220 μl of cold 1M Tris-HCl pH 7.4 (20mM final concentration) was added and the samples were centrifuged for 20 min at full speed (16000 rpm) at 4°C. The supernatant was rotated with 50 μl MagReSyn streptavidin beads (MR-STV0101, 2B Scientific), which were equilibrated with lysis buffer, at 4°C overnight. Samples were transferred into 1.5 ml centrifuge tubes and placed into a magnetic rack. Beads were washed twice with 1 ml wash buffer 1 (2% [wt/vol] SDS], once with 1 ml wash buffer 2 (0.1% [wt/vol] deoxycholate, 1% [wt/vol] Trition X100, 500 mM NaCl, 1 mM EDTA, 50 mM Hepes pH 7.4), and once with 1 ml wash buffer 3 (250 mM LiCl2, 0.5% [wt/vol] NP-40, 0.5% [wt/vol] deoxycholate, 1 mM EDTA, 10 mM Tris pH 8.1). Each washing step was performed for 8 min at RT using rotation. Proteins were eluted in 100 μl 2x reducing sample buffer (2% [wt/vol] SDS, 12% [wt/vol] sucrose, 0.004% [wt/vol] bromphenol blue, 50 mM Tris/HCl pH 6.8, 10% [wt/vol] 2-mercaptoethanol (or DTT) and 10 mM biotin) for 10 min at 90°C with mixing every 2 min. Biotinylation of proteins was confirmed using Western blotting and samples were analysed using liquid chromatography-tandem MS (LC-MS/MS).

To prepare sample for MS, 20 μl of eluted proteins were briefly run on SDS-PAGE (4 min at 160 V, 10% SDS gel [NP0301, Invitrogen]), stained with SimplyBlue SafeStain (LC6065, Invitrogen) for 1 hour and washed with ddH2O four times for 5 min each. For protein digestion, bands of interest were excised from the gel and dehydrated using acetonitrile followed by vacuum centrifugation. Dried gel pieces were reduced with 10 mM dithiothreitol and alkylated with 55 mM iodoacetamide. Gel pieces were then washed alternately with 25 mM ammonium bicarbonate followed by acetonitrile. This was repeated, and the gel pieces dried by vacuum centrifugation. Samples were digested with trypsin overnight at 37°C.

### Mass spectrometry data acquisition

Digested samples were analysed by LC-MS/MS using an UltiMate^®^ 3000 Rapid Separation LC (RSLC, Dionex Corporation, Sunnyvale, CA) coupled to an Orbitrap Exploris 480 (Thermo Fisher Scientific, Waltham, MA) mass spectrometer. Mobile phase A was 0.1% formic acid in water and mobile phase B was 0.1% formic acid in acetonitrile and the column used was a 250 mm x 75 μm i.d. 1.7 μM nanoE MZ PST CSH130 C18, analytical column (Waters). A 2 μl aliquot of the sample was transferred to a 5 μl loop and loaded on to the column at a flow of 300nl/min for 8 min, which then ramped to 5% B in 2 min. Peptides were separated using a gradient that went from 5% to 21% B in 44 min, then from 21% to 31% B in 7 min and finally from 31% B to 65% B in 1 min. The column was washed at 65% B for 4 min before dropping to 1% B in 1 min, and re-equilibration for a further 8 min. Data was acquired in a data dependent manner using a fixed cycle time of 1.5 sec, an expected peak width of 15 sec and a default charge state of 2. Full MS data was acquired in positive mode over a scan range of 300 to 1750 Th, with a resolution of 120,000, a normalised AGC target of 300% and a max fill time of 25 mS for a single microscan. Fragmentation data was obtained from signals with a charge state of +2, +3 or +4 and an intensity over 5,000 and they were dynamically excluded from further analysis for a period of 15 sec after a single acquisition within a 10ppm window. Fragmentation spectra were acquired with a resolution of 15,000 with a normalised collision energy of 30%, a normalised AGC target of 300%, first mass of 110 Th and a max fill time of 25 mS for a single microscan. All data was collected in profile mode.

The mass spectrometry proteomics data have been deposited to the ProteomeXchange Consortium via the PRIDE (Perez-Riverol et al., 2022) partner repository with the dataset identifier PXD037933.

All raw data were processed using MaxQuant software (v 1.6.10.43) (Table S2 and S3, Tyanova et al., 2016). Spectra were searched against the canine (Canis lupus familiaris) proteome obtained from Uniprot (November 2020) (UniProt, 2021). Biotinylation of lysine, methionine oxidation and N-terminal acetylation were set a variable modification, with carbamidomethylation of cysteine as a fixed modification. Precursor tolerance was set at 20 ppm and 4.5 ppm, for first and main search respectively, with MS/MS tolerance set at 20 ppm and up to two missed cleavages were allowed. The false discovery rate (FDR) of PSM and protein were set at 0.01 and “Match between runs” was enabled.

### Bioinformatical analyses

Protein intensities were exported from MaxQuant, normalised by median-centering and analysed through SAINTexpress software (v3.6.3) (Teo et al., 2014). Bait samples were grouped and analysed against their respective condition specific control samples. Significant proximal bait-prey interactions were taken using a Bayesian false discovery rate (BFDR) threshold of 0.05. Before any Gene Ontology (GO) analysis, human orthologs were obtained, where possible, for all quantified canine proteins using the Ensembl BioMart Service (Ensembl release 102) (Kinsella et al., 2011; Yates et al., 2020) and manual curation using the PANTHER database (v 16) (Mi et al., 2021). For seven protein hits we identified protein groups which cannot be distinguished with the current MS methodology. These hits comprised HLA-F/HLA-A/HLA-B/HLA-C/, RAB11A/RAB11B, ANP32D/ANP32A, TUBB2B/TUBB2A, HSPA1B/HSPA1A, DYNLRB2/MAP1LC3A, DNAJB7/DNAJB6, which were included separately for network and GO analyses. All subsequent GO functional analyses were then performed in the R environment (R Core Team, 2018) using the R package ClusterProfiler (v4.0.3) (Wu et al., 2021) against human GO annotations. For hierarchical clustering a list of prey proteins was taken that were significant at BFDR 0.05 for any bait in either condition. Hierarchical clustering was then carried out on the list of preys based on the Jaccard distance of significant (BFDR 0.05) and non-significant (BFDR > 0.05) interactions across all baits and both conditions. A heatmap was rendered to visualise the prey clusters, overlaying the log2 fold-change enrichment over BirA-myc control in each bait-condition. Network visualisation and analysis were performed using Cytoscape (v 3.9.0) (Su et al., 2014). Network analysis was performed using the stringApp plugin in cytoscape which is based on the STRING (search tool for the retrieval of interacting genes/proteins) database (v 11.5 accessed October 2021) (Doncheva et al., 2019; Szklarczyk et al., 2021). STRING scores of 0.7 and < 0.7 were distinguished and edges were discretely mapped accordingly. Protein-protein interactions were compared with the Biological General Repository for Interaction Datasets for DSC2, PKP2 and JUP (BioGRID v 4.4) (Oughtred et al., 2021) (available from https://thebiogrid.org/). Protein functions in Fig. 5 were annotated manually based on the primary literature and the Human Protein Atlas (Thul et al., 2017, available from http://www.proteinatlas.org). Graphing were performed using GraphPad Prism (v 9.2.0) (GraphPad Software, San Diego, California USA, www.graphpad.com). Venn diagrams were generated with the help of BioVenn (Hulsen et al., 2008). Figures were assembled in Adobe Illustrator (v 26.0.1).

## Supplemental Figures

**Figure S1.**
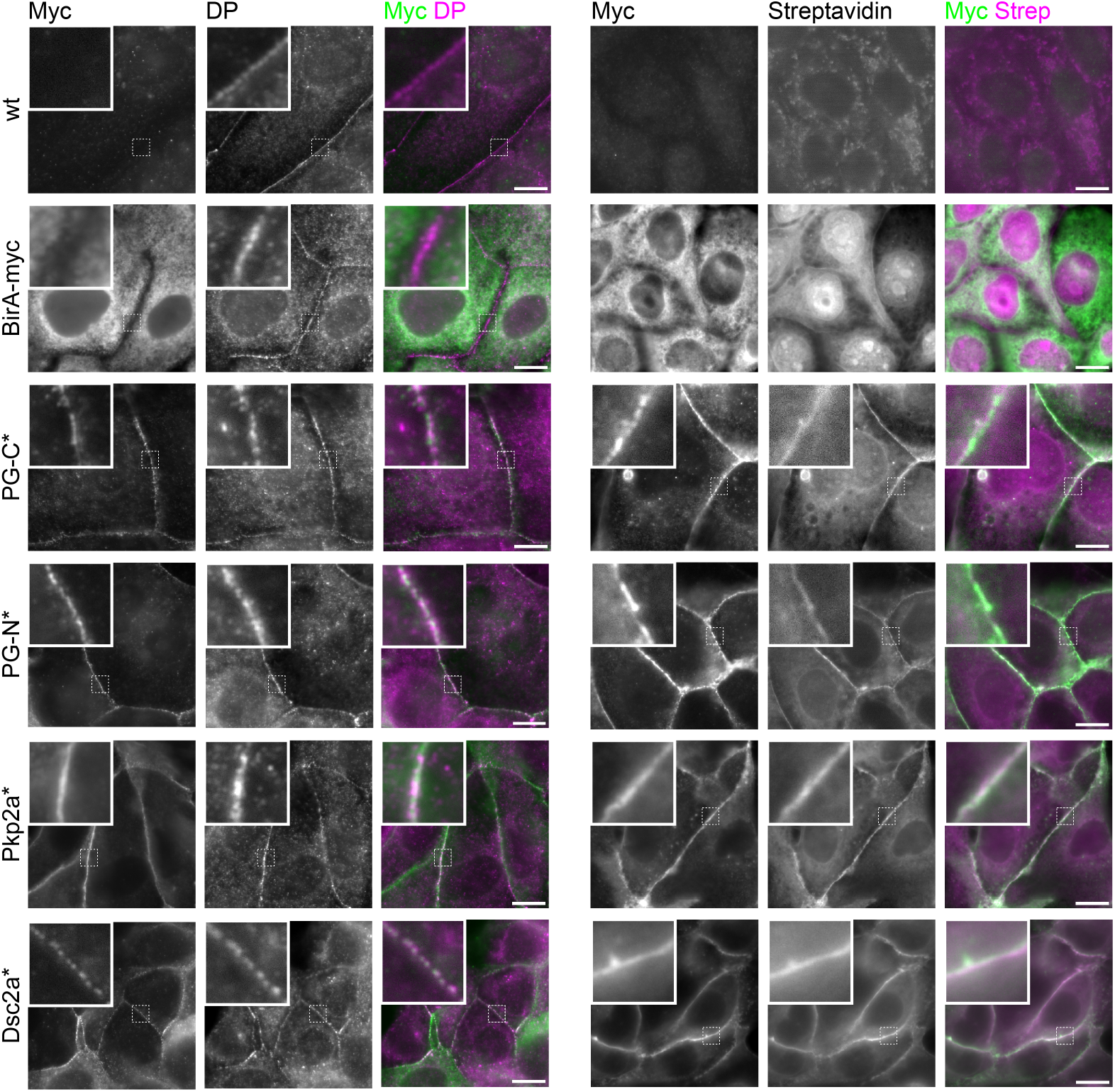
BioID constructs to generate desmosome proximitome localise to desmosomes. Confocal microscopy images of MDCK cells stably expressing BioID constructs as shown in Figure 1 and the parental wild-type cells. Cells were cultured subconfluent for 24 h including 16 h of incubation with 100 μM biotin before immuno-labelling desmoplakin (DP) (left panel) or for myc and fluorescently conjugated streptavidin (right panel). Data representative of three biological repeats. Scale bars: 10 μm.

**Figure S2.**
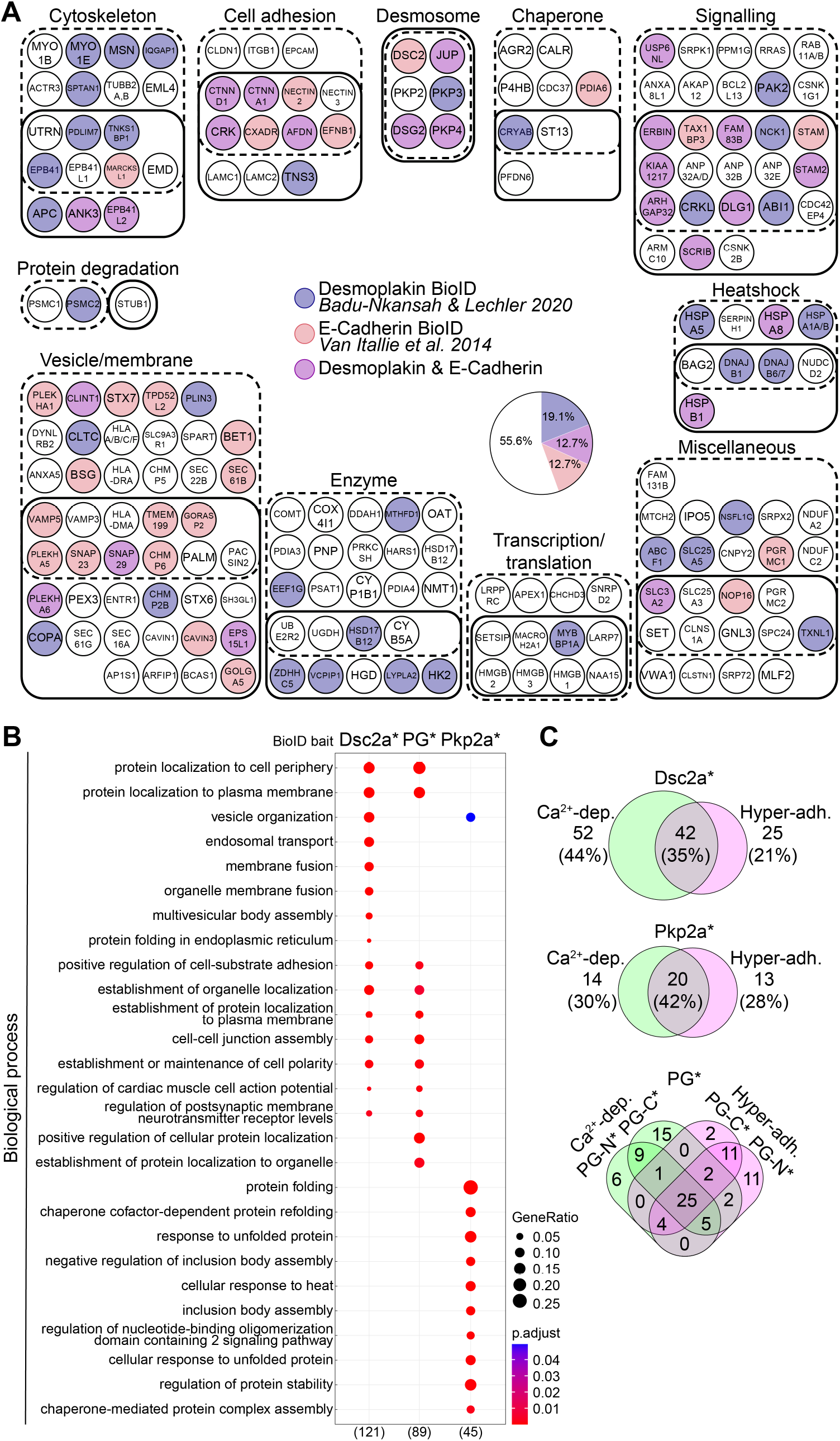
Further analysis of the desmosome proximitome. Accompanies figure 2 and 3. **(A)** Functional network of desmosome BioID data presented in this study (combined proximal proteins to desmocollin 2a-BirA-myc (Dsc2a*), myc-BirA-plakoglobin (PG-N*), plakoglobin-BirA-myc (PG-C*), myc-BirA-plakophilin 2a (Pkp2a*); BFDR ≤ 0.05) compared to BioID studies of desmoplakin and E-cadherin (Badu-Nkansah and Lechler, 2020; Van Itallie et al., 2014). Proteins were annotated using the primary literature and the Human Protein Atlas and classified into the indicated categories (see text relating to figure 5). Boundaries indicate whether the prey proteins were significantly enriched in data curated from MDCK cells cultured confluently for either 1 day and thus with Ca^2+^-dependent desmosomes (dotted line) or for 5 days and thus hyper-adhesive desmosomes (solid line). Proximal prey proteins are coloured in purple when they were also identified in the desmoplakin interactome, pink when they were present in the E-cadherin interactome and in magenta when they were present in both the desmoplakin and E-cadherin datasets. Pie chart shows the overlapping prey proteins identified in this study in percentage. **(B)** GO enrichment analysis of the 189 proteins partial desmosomal proximitome. The top 10 overrepresented terms of each bait (Dsc2a*, Pkp2a* and PG* [PG-N* and PG-C* combined]) under the biological process category are shown. The number of annotated proteins is shown in brackets. (Note 7 protein hits encompassed proteins groups of which all members were included for GO analysis because we could not distinguish between different isoforms. For details see Materials and Methods.) p.adjust, adjusted P value. GeneRatio, proportion of total proteins identified in each GO term. **(C)** Venn diagrams illustrating the number of overlapping proteins depending on the bait and adhesion state.

**Figure S3.**
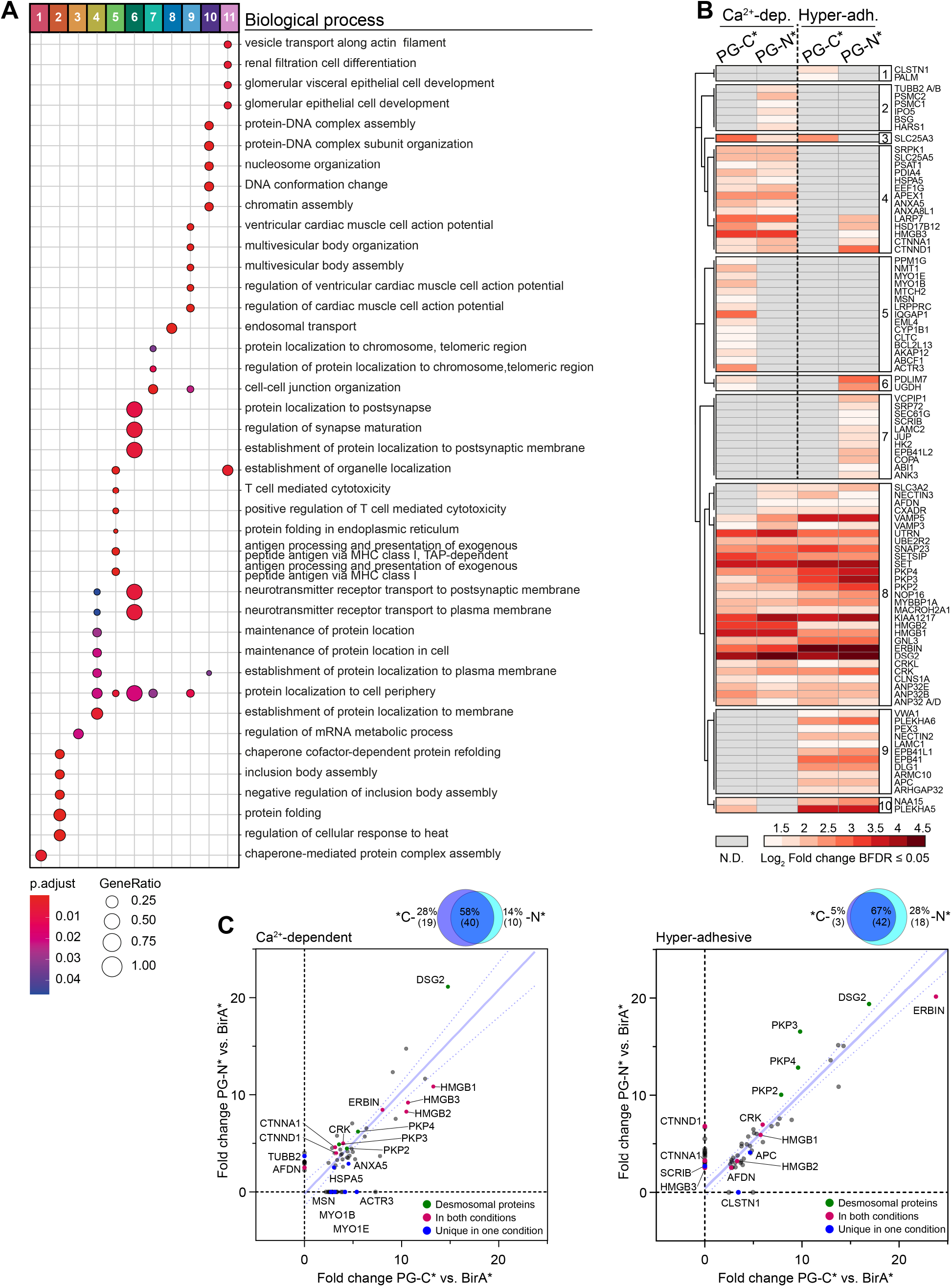
Functional enrichment analysis of prey clusters and hierarchical clustering of preys of plakoglobin. Accompanies figure 4. **(A)** GO analysis of the prey clusters identified from hierarchical clustering of the desmosomal proximitome (Fig. 4 A). The top ten overrepresented GO terms under the biological process category are shown. p.adjust, adjusted P value. GeneRatio, proportion of total proteins identified in each GO term. **(B)** Hierarchical clustering was performed on the proteins identified in the plakoglobin (PG) BioID proximitome and the results are displayed as a heatmap. **(C)** Scatter plot and area-proportional Venn diagram showing the relationship of PG-N* and PG-C* prey proteins in either Ca^2+^-dependent or hyper-adhesive conditions (fold change enrichment over BirA*). The 95% confidence interval of the regression line is displayed as dotted confidence band.

**Figure S4.**
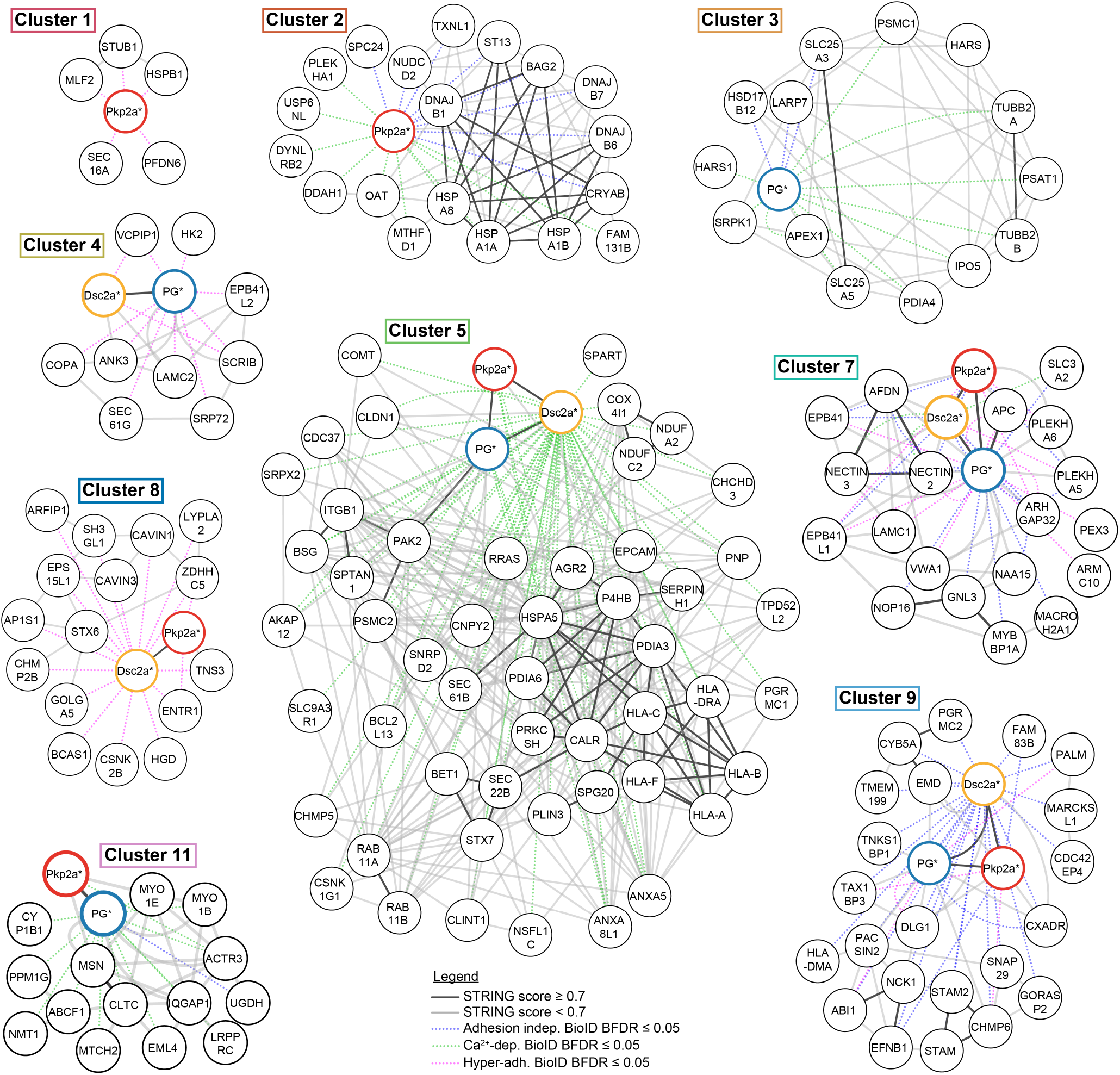
Network analysis of prey clusters. STRING network analysis of the prey clusters identified from hierarchical clustering of the desmosomal proximitome (Fig. 4). Nodes of bait proteins are colour codes with Pkp2a* in red, Dsc2a* in yellow and PG* (merged PG-N* and PG-C*) in blue. Edges indicate protein-protein interactions: solid dark grey lines indicate a STRING score ≥ 0.07; solid light grey lines a STRING score below 0.7 and dotted lines BioID proximity with a BDFR ≤ 0.05 presented in this study (blue = adhesion independent, green = Ca^2+^-dependent, magenta = hyper-adhesive).

